# Heterologous expression of pyruvate formate lyase enhances cell growth of *Clostridium ljungdahlii* during microbial electrosynthesis

**DOI:** 10.1101/2025.09.12.675768

**Authors:** Chaeho Im, Adolf Krige, Kaspar Valgepea, Oskar Modin, Yvonne Nygård, Carl Johan Franzén

**Affiliations:** Division of Industrial Biotechnology, Department of Life Sciences, Chalmers University of Technology, Göteborg, Sweden; Institute of Bioengineering, University of Tartu, Tartu, Estonia; Division of Water Environment Technology, Department of Architecture and Civil Engineering, Chalmers University of Technology, Göteborg, Sweden; VTT Technical Research Centre of Finland, Espoo, Finland

**Keywords:** Bioelectrochemical system, microbial electrosynthesis, *Clostridium ljungdahlii*, acetogen, pyruvate formate lyase, formate

## Abstract

Slow cell growth and low biomass yields are hurdles for gas fermentation by acetogens. Microbial electrosynthesis (MES) utilizes acetogens as biocatalysts to reduce CO_2_ and produce commodity chemicals using electricity. However, limited electron supply in a bioelectrochemical system (BES) aggravates the poor cell growth of acetogens resulting in low productivities. Formate is a soluble C1 feedstock that can be produced by CO_2_ reduction (1). Thus, assimilation of formate can unburden the amount of electrons required for MES. *Acetobacterium woodii* posseses multiple pyruvate formate lyase (PFL) genes that are upregulated during formatotrophic growth. In this study, *Clostridium ljugdahlii* was engineered to heterologously express a *pfl* geneset of *A. woodii* to increase the cell growth of *C. ljungdahlii* during microbial electrosynthesis. Different combinations of *pfl A* and *pfl B* from *A. woodii* were tested in *C. ljungdahlii* to find the best working combination under control of *P_fdx_*-riboswitch expression system. Expression of *pfl* B1 and *pfl*A of *A. woodii* showed higher maximum OD of *C. ljugdahlii* under H_2_:CO_2_ condition with supplementation of 80 mM sodium formate. More than 40 mM of sodium formate caused significantly longer lag phase but the lag phase could be shortened after adaptation in 80 mM of sodium formate. At the end, the engineered strain showed improved cell growth (OD_max_ 0.22 ± 0.05) and acetate production (21.8 ± 5.6 mM) during microbial electrosynthesis compared to the control strain (OD_max_ 0.10 ±0.06 and 10.2 ±2.5 mM acetate). These results will be useful for strain development for microbial electrosynthesis, as well as gas fermentation.

**Highlights:**

- Different combinations of pyruvate formate lyases (*pfl B*s) and pyruvate formate lyase acting enzymes (*pfl A*s) from *Acetobacterium woodii* were tested to improve the cell growth of *Clostridium ljungdahlii* during gas fermentation and microbial electrosynthesis
- Heterologous expression of *pfl B1* and *pfl A* using riboswitch expression system improved cell growth of *C. ljungdahlii* under H_2_:CO_2_ condition, even without inducer and formate supplementation.
- High concentraton of sodium formate caused longer lag phase, which was shortened by adaptation when *pfl* from *A. woodii* was heterologously expressed.
- Cell growth and acetate production of the engineered *C. ljungdahlii* strain improved during microbial electrosynthesis

## 1. Introduction

Acetogens fix CO_2_ via the Wood-Ljungdahl Pathway to synthezise acetyl-CoA. Acetyl-CoA can be converted further into pyruvate to provide the cell with building blocks for biosynthesis, acetate for ATP synthesis, and other metabolic products like acetaldehyde and ethanol for redox balancing (2). The greenhouse gas CO_2_ is a representative residual carbon emitted from many industries, and CO_2_ emission should be reduced to mitigate climate change by minimizing the greenhouse effect. Autotrophic growth of acetogens gives a promising application of acetogens, namely gas fermentation leading to CO_2_ capture and utilization via the Wood-Ljungdahl Pathway. However, low ATP yield from the metabolism of CO_2_ and H_2_ and low solubility of H_2_ as an electron donor for CO_2_ reduction result in slow cell growth and low maximum cell density during gas fermentation.

Bioelectrochemical systems (BESs) utilizes the interactions between electro-active bacteria and electrodes for the regulation of electro-active bacterial metabolism. Microbial electrosynthesis (MES) is a BES used to valorize residual carbons and produce value-added chemicals by using electricity to supply electro-active bacteria with reducing equivalents.

Electro-active acetogens can be used as biocatalysts in MES to valorize CO_2_ and produce commodity chemicals. *Sporomusa ovata*, *Sporomusa spheroides*, *Sporomusa silacetica*, *Clostridium aceticum*, *Clostridium ljungdahlii*, *Clostridium scatologenes*, *Acetobacterium woodii*, and *Moorella thermoacetica* have showen their ability to produce acetate from CO_2_ in a BES (3–6). Among electro-active acetogens tested, *C. ljungdahlii* is an industrially relevant acetogen strain with the ability to utilize both CO_2_ and CO and produce ethanol. *C. autoethanogenum* and *C. ljungdahlii* are genetically closely related strains, and *C. autoethanogenum* is already being used for commercialized gas fermentation processes (8, 9). While pure *C. autoethanogenum* culture has not shown its electro-activity in a BES, *C. ljungdahlii* has been tested in a BES several times (3, 5, 10, 11). Therefore, understanding and engineering microbial electro-synthesis from CO_2_ using *C. ljungdahlii* for improved performance can benefit valorizing gaseous waste in a BES.

BES studies using acetogens belonging to the *Clostridium* genus were mostly done using *C. ljungdahlii* because of its industrial relevance and the opportunity of strain engineering. *C. ljungdahlii* was first tested in a BES in a screen for acetogens capable of MES (5). Around the same time, *C. ljungdahlii* was metabolically engineered for the first time for alcohol production (2, 12). Later, *C. ljungdahlii* was engineered to heterologously express formate dehydrogenase from *Escherichia coli*, and the engineered *C. ljungdahlii* showed improved electricity generation in a microbial fuel cell with elevated [NADH]/[NAD^+^] level (13). Electrochemically-produced H_2_-driven MES using *C. ljungdahlii* was demonstrated by using an assembly of graphite felt and stainless steel mesh as a cathode (11). The cathode material decreased the overpotential and improved volumetric acetate production rates up to 60-70 times higher at – 0.9 V (vs Ag/AgCl) cathode potential (CP) than observed in the first MES study using *C. ljungdahlii* at – 0.6 V CP with a graphite stick as a cathode material (11). The study showed the relevance of the H_2_ route for CO_2_ reduction and planktonic cells during MES (11). Boto et al., demonstrated how to control planktonic and biofilm cell growth of *C. ljungdahlii* during MES and stressed the importance of H_2_-mediated MES using planktonic *C. ljungdahlii* for a robust performance (10). Nonetheless, the maximum optical density (OD) achieved in the study was around OD 0.36, which is much lower than in typical [CO_2_:H_2_]=[20:80] gas fermentation (∼OD 1.0), probably due to limited electron supply (10, 14).

The maximum OD of *C. ljungdahlii* is lower when H_2_ is used as the electron donor compared to CO (15). This is because reduced ferredoxin produced from H_2_ oxidation must be utilized for CO_2_ reduction to CO for acetyl-CoA synthesis in the Wood-Ljungdahl pathway, which can be compensated by supplementation of small amount of CO (16). Reduced ferredoxin is an important electron carrier in acetogens as it is required in several processes, e.g., ATP synthesis *via* the Rnf complex route, ethanol production via aldehyde:ferredoxin oxidoreductase, the Wood-Ljungdahl pathway, Nfn complex, and pyruvate synthesis (17). To achieve higher biomass during MES, carbon flux towards gluconeogenesis should be increased (18). Pyruvate synthesis from acetyl-CoA is the first step in gluconeogenesis for producing cellular building blocks, providing precursors for amino acids and nucleotide synthesis essental for cell growth (19).

There are two ways of producing pyruvate for cell growth from Acetyl-CoA in acetogens: one is pyruvate:ferredoxin oxidoreductase (PFOR) and the other one is pyruvate formate-lyase (PFL) (Fig. 1A) (23). PFOR requires reduced ferredoxin and CO_2_, while PFL requires formate to produce pyruvate from acetyl-CoA (2, 24). Therefore, utilizing formate for pyruvate formation through the PFL route could bypass the use of reduced ferredoxin for the ATP synthesis required for cell growth. Formate can be produced electrochemically and biologically (1). Formate is an emerging C1 feedstock and important intermediate in acetogens, serving as a carbon and electron source (1, 25). While some acetogens can grow directly on formate (23, 26), formate can also be used as a co-substrate during autotrophic growth (27).

In this study, *C. ljungdahlii* was engineered to heterologously express PFL from *A. woodii* to be able to grow better during MES using formate as a co-substrate. Wild-type and engineered *C. ljungdahlii* strains were tested in serum bottles with different sodium formate concentrations under H_2_:CO_2_ conditions, as well as in a BES. The results of this study will contribute to push the barrier of gas fermentation towards better performance.

**Figure 1.**
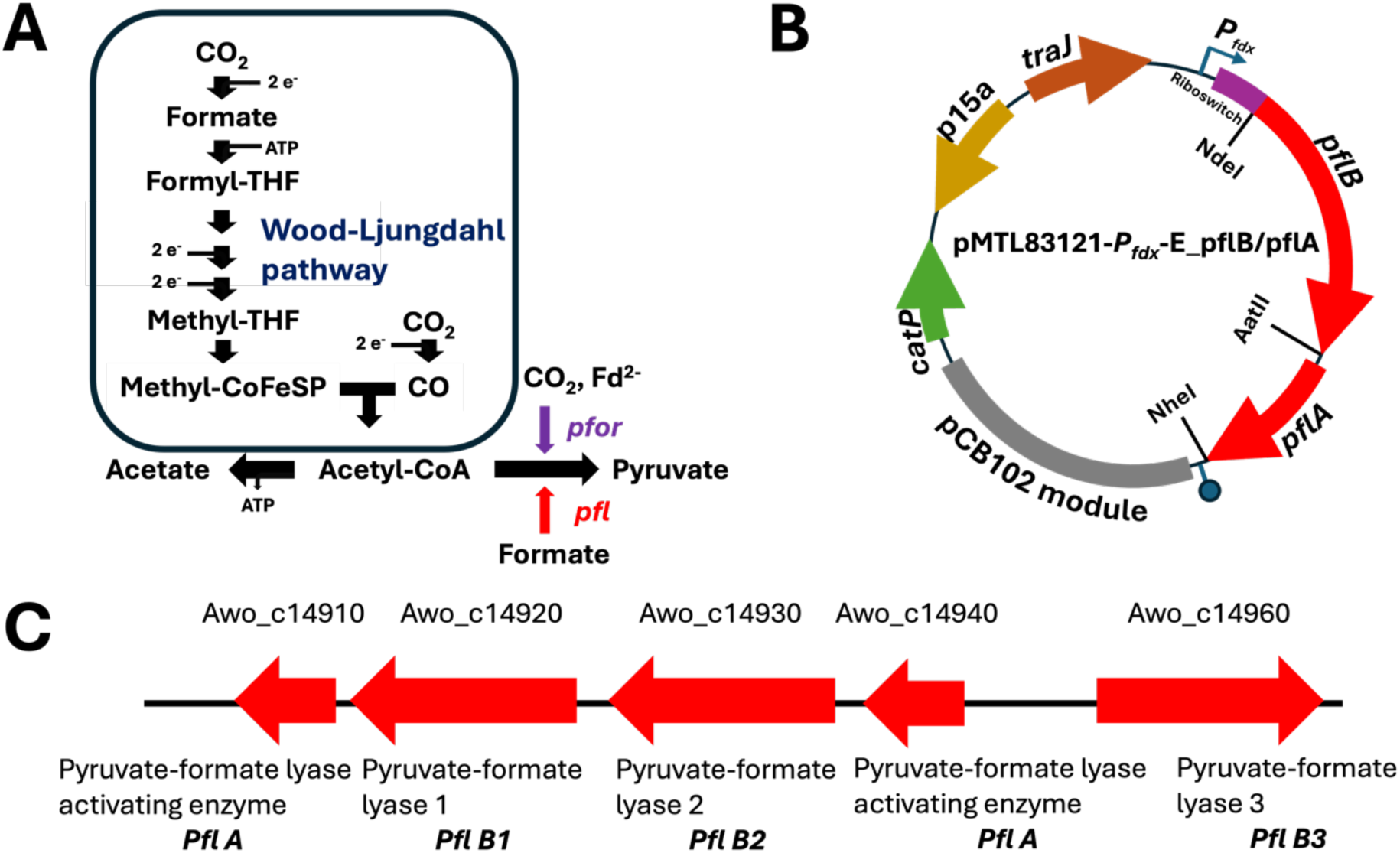
(A) Schematic diagram of the Wood-Ljungdahl metabolic pathway with acetate and pyruvate production routes. THF: tetrahydrofolate, pfor: pyruvate:ferridoxin oxidoreductase, pfl: pyruvate formate lyase. (B) Plasmid map of pMTL83121-*P_fdx_*_E_pflB/pflA. (C) pyruvate formate lyase and pyruvate formate lyase activating enzyme genes in A. woodii genome

## 2. Materials and Methods

### 2.1 Bacterial strians and growth conditions

*Clostridium ljungdahlii* DSM13528 was purchased from Deutsche Sammlung von Mikroorganismen und Zellkulturen (DSMZ) GmbH (Braunschweig, Germany). *Escherichia coli* NEB 10 beta was purchased from New England Biolabs (Massachusetts, USA) and employed for plasmid propagation, cloning, and conjugation. *E. coli* strains were cultivated at 30 °C in LB or TB medium in the presence of antibiotics if necessary (25 µg/mL chloramphenicol, 50 µg/mL kamamycine). The bacterial cultures were stored in 20 % glycerol in cryotubes at – 80 °C. Bacterial strains used in this study are listed in Table 1.

*C. ljungdahlii* was cultivated in 20 mL of anaerobically prepared modified DSMZ 879 medium under 2 bars of over-pressurized gas [H_2_:CO_2_]=[80:20] in 200 mL rubber stopper sealed serum bottles, with 5 µg/mL thiamphenicol when necessary (3). The cultures were placed horizontally in a temperature-controlled shaking incubator at 37 °C and 160 rpm. The headspace of the serum bottles was re-filled with the same gas when the headspace pressure dropped to less than 1 bar over-pressure.

### 2.2 DNA manipulation

Genomic DNA of *A. woodii* was extracted using MasterPure^TM^ Gram-positive DNA purification kit (LGC Bioresearch Technologies, Hoddesdon, UK) for polymerase chain reactions (PCRs). PCRs were carried out using Q5 High-fidelity DNA polymerase (New England Biolabs, Massachusetts, USA). All primers for PCRs were synthesized by Eurofins (Luxembourg) and are listed in Table 1. Restriction enzymes and T4 DNA ligase were purchased from New England Biolabs. Plasmids extraction was done using GeneJET Plasmid Miniprep Kit (Thermo Fisher Scientific, Massachusetts, USA). Sanger sequencing of plasmids and amplicons was carried out by Eurofins. All plasmids used in this study are derived from the pMTL80000 series of modular *E.coli*-*Clostridium* shuttle vector and are listed in Table 1 (28).

pMTL83121-empty was constructed as the backbone plasmid from a modular *E. coli* – *Clostridium* shuttle vector system for this work (28). P*_fdx_*-riboswitch E was amplified using PCR with primer Pfdx.ribo_F and Pfdx.ribo_R using pRECas1 as the template. P_fdx_-riboswitch E amplicon and pMTL83121 were double-digested using NotI and NdeI and ligated to create pMTL83121-*Pfdx-E*_*empty*. *A. woodii pfl B1*, *B2*, *B3*, *A1*, and *A2* were amplified using PCR, and the primers listed in Table 1. *A. woodii pfl B1*, *B2*, and *B3* were double-digested using NdeI and AatII, and *A. woodii pfl A1* and *A2* using AatII and NheI. pMTL83121-*Pfdx-E*_*empty* was double-digested using NdeI and NheI. All products from double-digestion using restriction enzymes were purified using gel electrophorasis and GeneJet Gel Extraction Kit (Thermo Fisher Scientific, Massachusetts, USA). The purified products were ligated using T4 ligase with different combinations to create the plasmids listed in Table 1.

**Table 1.**
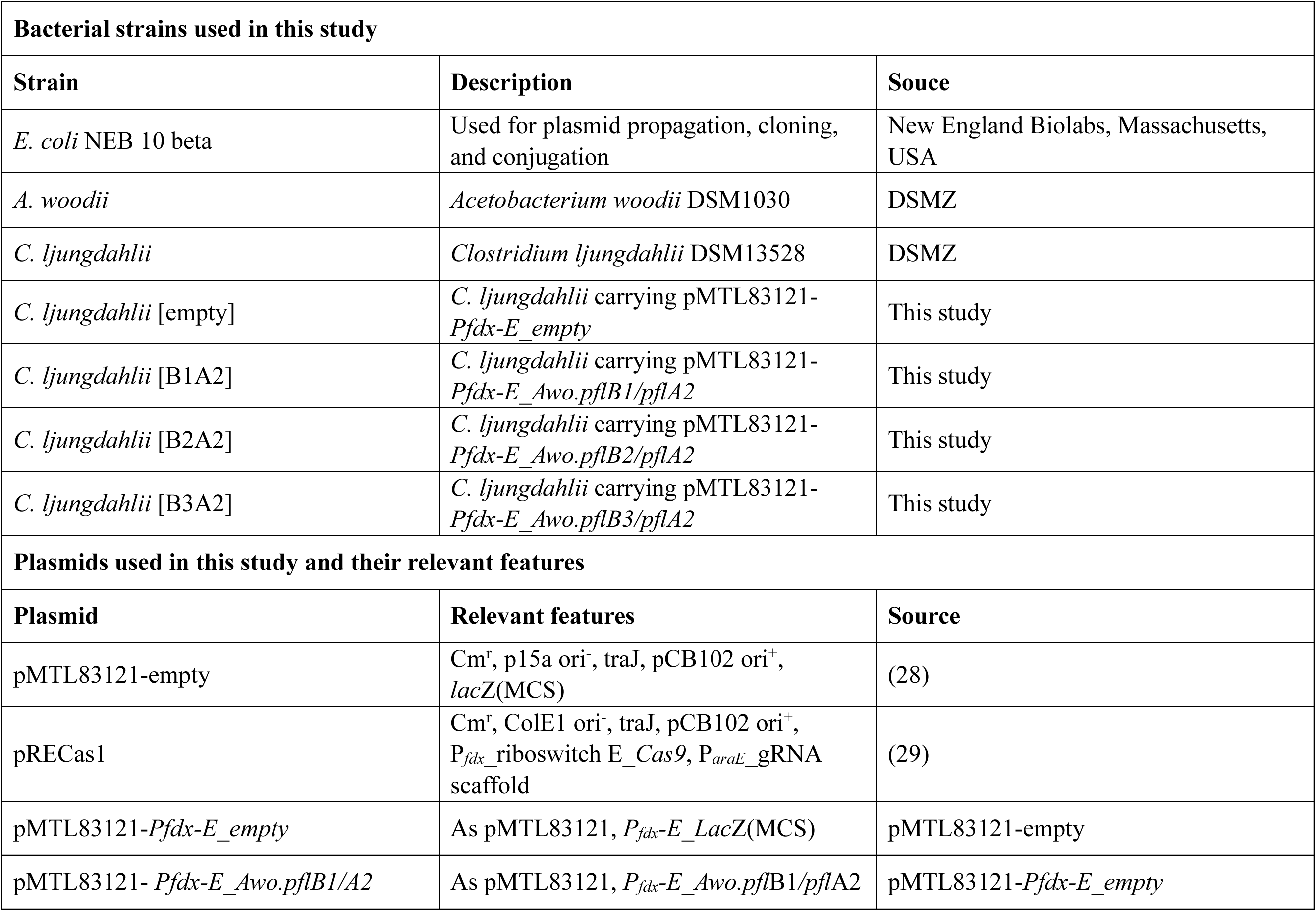
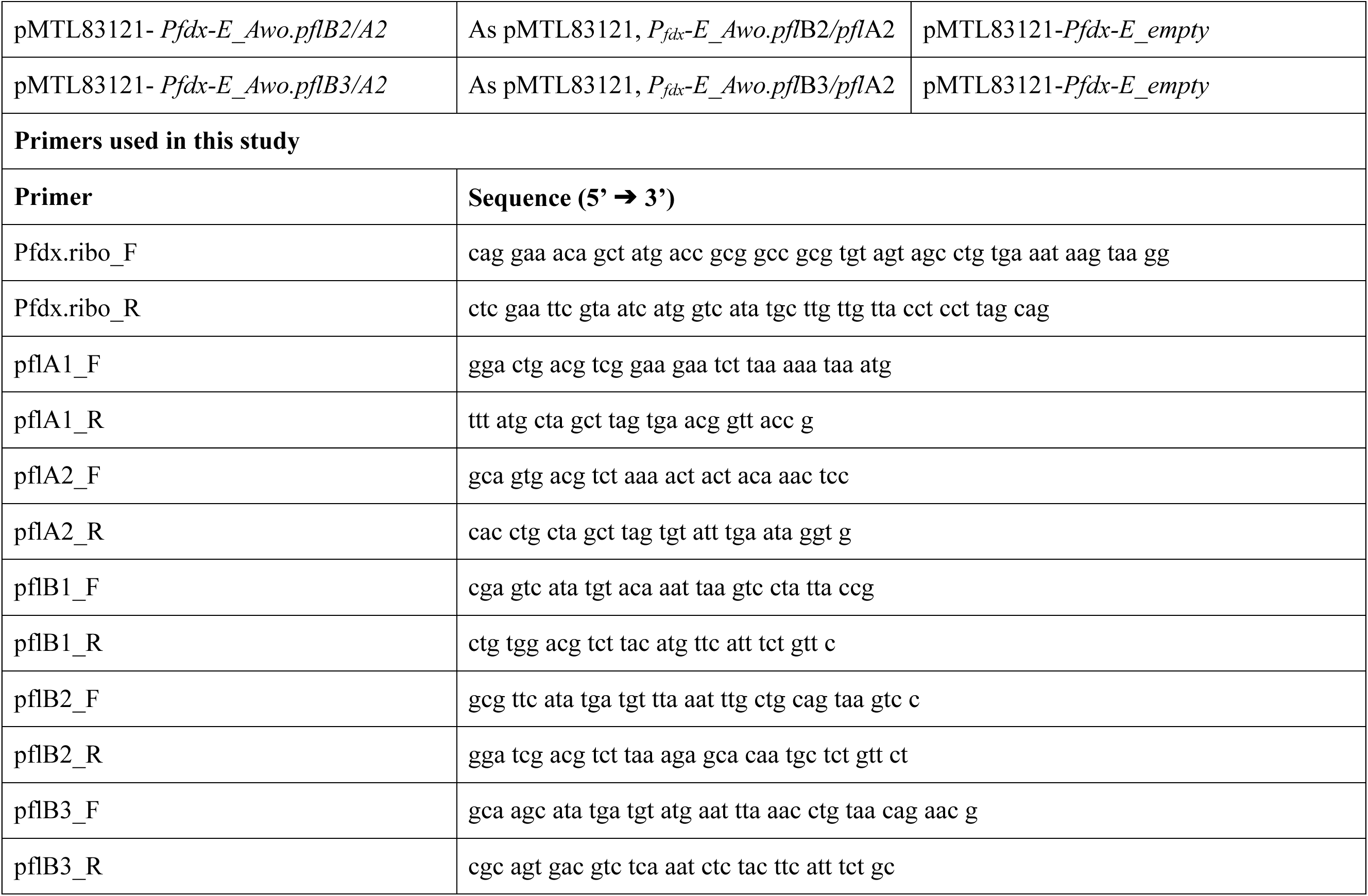
Bacterial strains, plamids, and primers used in this study.

### 2.3 Conjugal plasmid transfer into *C. ljungdahlii*

The general procedure of conjugal plasmid transfer was adapted from Woods et al. (30). Briefly, Plasmids were transfomed into *E. coli* NEB 10 beta with pRK2013 helper plasmid. *E. coli* NEB 10 beta carring each plasmids with pRK2013 helper plasmid was cultivated overnight in LB with 25 µg/mL chloramphenicol and 50 µg/mL kanamycin. Next day, 1 mL of the overnight cultures was harvested in 1.5 mL Eppendorf tubes and centrifuged at 4000 x g for 2 minutes. The supernatants were discarded and the cell pellets were rinsed gently with phosphate-buffered saline solution (pH 6.0), twice. The cell pellets were resuspended gently in 200 µL of wild-type *C. ljungdahlii* culture grown overnight in YTF (Yeast extract-Tryptophan-Fructose) medium. Spots of 25 µL of the recipient *C. ljungdahlii* and donor *E. coli* strain culture-mix were dispensed on YTF agar plates and were left to dry for spot-mating. After 16-20 hours of spotting on the agar plates, the dried cell pellets on the agar plate were scraped off by a inoculation loop and resuspended in 400 µL of YTF liquid medium. Different dilutions of the resuspended culture were spread on YTF agar plates containing 5 µg/mL thiamphenicol and 500 µg/mL D-cycloserine. When colonies were observed, a single colony was re-streaked on a YTF agar plate containing 5 µg/mL thiamphenicol and 500 µg/mL D-cycloserine to avoid carry-over of the donor *E. coli* strain. The plasmid transformation was double-checked by plasmid extraction and re-transformation into *E. coli* strain (31).

### 2.4 Bioelectrochemical reactor operation

H-type BES reactors (Adams & Chittenden Scientific Glass Coop, Berkeley, CA, USA) were used for MES (3). The cathode and anode chambers of the BES were connected by an NW40 high-temperature chain clamp (Evac AG, Grabs, Switzerland) and separated by an Aquivion® E98-05S membrane (d = 4 cm; Solvey, Brussels, Belgium). Each chamber contained 250 mL modified DSMZ879 medium (pH 5.0) without resazurin. A graphite felt (2.5 cm × 4.0 cm; Alfa Aesar, Massachusetts, USA) connected with a titanium wire (Alfa Aesar, Massachusetts, USA) was used as cathode, while a platinised titanium wire (MAGNETO special anodes B. V., Schiedam, The Netherlands) was used as anode. Cells from the preculture were inoculated to an OD_600_ of 0.05. The cultures were sparged continuously with 100 % CO_2_ at 10 mL/min. The current was maintained using a KP 07 potentiostat/galvanostat (Bank Elektronik - Intelligent Controls GmbH, Pohlheim, Germany). 1 mL of sample was taken from both the cathode and anode chambers every 24 h to determine OD_600_, pH, and metabolite composition. Each BES experiment was performed in duplicate.

### 2.5 Analytical chemistry

Optical density at 600 nm (OD_600_) was measured using a WPA S1200+ visible spectrophotometer (Biochrom, Cambridge, UK). A pH meter (Mettler Toldedo, Columbus, OH, USA) was used to measure the pH of samples. Metabolite analysis was performed using a high-performance liquid chromatography system equipped with a refractive index detector (at 40 °C) and a UV detector (210 nm) (Jasco, Tokyo, Japan), as well as a ROA-Organic acid H^+^ (8%) column (Rezex, Torrance, CA, USA). The oven temperature was maintained at 60 °C and the column was eluted with 5 mM H_2_SO_4_ at 0.6 mL/min. The collected liquid samples were centrifuged at 12,000 rpm for 5 min, followed by filtration through a 0.2 -μm nylon syringe filter before sample injection in the HPLC.

## 3. Results and Discussion

### 3.1 Sodium formate tolerance of *C. ljungdahlii*

We first tested tolerance of *C. ljungdahlii* to sodium formate to see what concentration of sodium formate should be used for the screening of the combination of *pfl B* and *pfl A*. When *C. ljungdahlii* was cultivated at initial pH 5.0, which has been tested for microbial electrosynthesis using *C. ljungdahlii* (3), the cell growth was only possible in up to 20 mM sodium formate (Table 2).

**Table 2.** Growth of *C. ljungdahlii* on different sodium formate concentrations.

|  | 0mM | 10 mM | 20 mM | 40 mM | 80 mM |
| --- | --- | --- | --- | --- | --- |
| pH 5.0 | + | + | + | - | - |
| pH 5.7 | + | + | + | + | + |

Cultivation of *C. ljungdahlii* at initial pH 5.7 enables the cell to grow 80 mM sodium formate, which is the highest sodium formate concentration tested in this study (Table 2). Therefore, 80 mM of sodium formate with initial pH 5.7 was chosen for the screening of the most effective *pfl B* and *pfl A* combination. A smaller addition of formate might not be enough for showing the effect of heterologous expression of PFL.

Formate inhibits cell growth of microorganims (32). This is because undissociated organic acids diffuse into cell membrane and increases internal proton concentration, which can damage DNA, affect DNA synthesis, and uncouple the proton motive force across the inner membrane for ATP synthesis (33). This becomes even more severe with more acidic pH as the ratio of undissociated to dissociated organic acid increases. Therefore, it is very likely that the formate tolerance of *C. ljungdahlii* was limited to 20 mM sodium formate at pH 5.0 while it could tolerate 80 mM sodium formate at pH 5.7 (Table 2).

### 3.2 Cloning of pyruvate formate lyase from *Acetobacterium woodii*

*Acetobacterium woodii* can grow solely on formate (23). Transcriptomic study using *A. woodii* grown on formate showed upregulation of PFL mRNA expression to produce pyruvate for gluconeogenesis using formate (23). *A. woodii* genome contains three *pfl B* genes: *pfl B1*, *B2*, and *B3* (Awo_c14920, Awo_c14930, and Awo_c14960, respectively) and two *pfl A* genes: *pfl A1* and *A2* (Awo_c14910 and Awo_c14940) (Fig. 1 C). In order to find out the most effective combination of *pfl B* and *A*, all the combinations were attempted to be cloned in pMTL83151 with *P_fdx_* promoter. However, only cloning of *pfl A* (Awo_c14940) alone was successful. Otherwise, cloning of each *pfl B* and *pfl A* (Awo_c14910) showed negative results or it was confirmed by Sanger sequencing that the start codon of the genes were mutated. Cloning of *pfl B* with *pfl A*(Awo_c14940) was possible only when low copy number replication of origin (p15a) and P_fdx_ promoter with Theophylline-riboswitch were used together (Fig. 1 B) (29, 34).

Restriction enzyme NdeI used for *pfl B* cloning contains a start codon ATG. The mutations observed in the start codon region of *pfl* while cloned in a high copy number plasmid and the strong promoter (*P_fdx_*) might be because of toxicity of the genes. Also, mRNA abundance of *pfl B* and *pfl A* in *A. woodii* under formatotrophic condition is low (23). Therefore, tightly regulated expression of PFL seems to be important.

A synthetic Theophylline riboswitch was used in *Clostridium* genus first for CRISPR/Cas9 system because of toxicity of uncontrolled Cas9 protein expression (29, 35). Theophylline-responsive riboswitch has been tested in various bacteria including several *Clostridium* species (29, 36–41). Theophylline riboswitches forms small hairpins to block both the ribosomal binding site (RBS) and the start codon (AUG) of mRNAs so that translation does not happen, while a conformational change in the presence of Theophylline allows translation by releasing RBS and the start codon (34). Therefore, it seems that Theophylline-responsive riboswitch expression system is controlled by thermodynamics rather than being species dependent (34). The disadvantage of a riboswitch expression system is that it is a translational inducible expression system which means it can control the expression of only one gene. To control more genes, the riboswitch must be placed in every RBS of each gene. Therefore, it should be noted here that *pfl A2* (Awo_c14940) was constitutively expressed while only *pfl B* was under the control of the riboswitch. Therefore, it would not be surprising that cloning *pfl A1* (Awo_c14910) were impossible, if overexpression of Awo_c14910 is toxic to the cells. On the other hand, *pfl B* activation requires *pfl A*. PFL is a glycyl radical enzyme requiring post-translational activation by the radical S-adenosylmethionine enzyme (*pfl A*) (42). The *pfl B* and *pfl A* association rate is very slow (42). Therefore, the constitutively overexpressed *pfl A2* (Awo_c14940) might increase the PFL reaction rate compared to if the *pfl A* expression also would be limited by the riboswitch.

### 3.3 Growth of *C. ljungdahlii* strains expressing *A. woodii pfl A* and *pfl B*

Wild-type and engineered strains were tested under H_2_:CO_2_ atmosphere plus 80 mM sodium formate to screen the most effective combination of *pfl B* working with *pfl A2* (Awo_c14940). Since the Theophylline riboswitch expression system combined with *P_fdx_* promoter was used, both presence and absence of 1 mM Theophylline as the translational inducer were tested. All the strains with and without Theophylline showed 4 - 6 days of lag phase under 80 mM sodium formate condition, except wild-type and *C. ljungdahlii* [B1A2] (*C. ljungdahlii* carrying pMTL83121-*Pfdx-E_Awo.pflB1/pflA2*) strains (Fig. 2).

**Figure 2.**
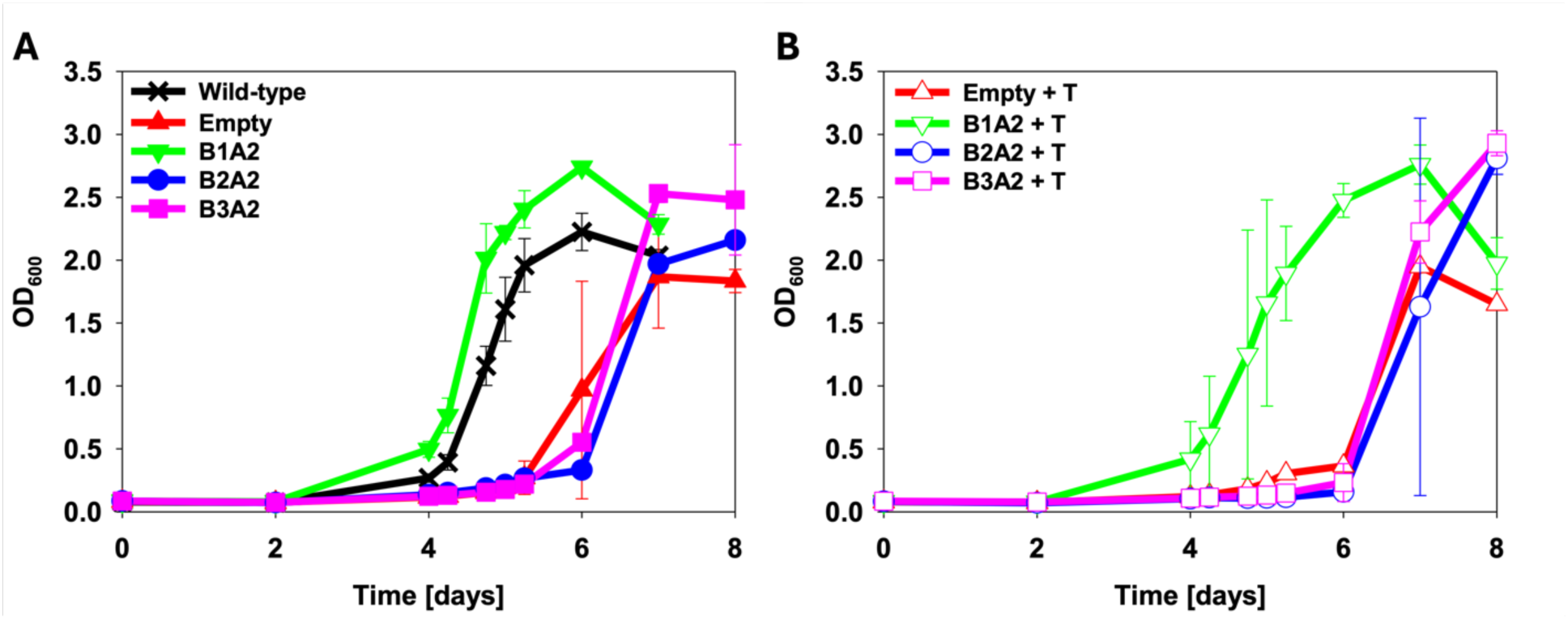
Time-dependent OD profile of C. ljungdahlii harboring pMTL83121-Pfdx-E with different pfl B and A combinations under H_2_:CO_2_ plus 80 mM sodium formate condition. A) without and B) with addition of 1 mM of Theophylline supplementation as translational inducer for the riboswitch. Filled symbols indicate without addition of the inducer, and open symboles indicate with addition of the inducer. Error bars indicate standard deviation (n=2, B2A2 and empty vector+T: n=1). See supplementary information (Fig. S1) for individual OD profiles.

The wild-type strain grew with the growth rate of *μ* = 0.078 ± 0.001 h^-1^ after 4 days of lag phase regardless of the presence or absence of the inducer. The control strain (*C. ljungdahlii* carrying pMTL83121-*P_fdx_-E*_empty) showed longer lag phase of about 5 days and growth rates of *μ* = 0.072 ± 0.014 h^-1^ without and *μ* = 0.070 h^-1^ with the inducer, respectively (Fig. 2). The lag phase was restored to 4 days with *pflB1* and *pflA2* expression. The specific growth rate was unaffected in the absense of Theophylline (*μ* = 0.078 h^-1^) but decreased to *μ* = 0.066 ± 0.004 h^-1^ in the presence of the inducer (Fig. 2). In all cases except *C. ljungdahlii* [B2A2] without the inducer (OD_max_ = 2.16), the maximum OD was increased compared to wild-type (OD_max_ = 2.23 ±0.15) and the negative control *C. ljungdahlii* [empty] (OD_max_ = 1.87 ±0.41 and 1.95, without and with the inducer, respectively) (Fig. 2). Even though the highest maximum OD was observed from *C. ljungdahlii* [B3A2] with the inducer (OD_max_ = 2.93 ± 0.1), *C. ljungdahlii* [B1A2] (OD_max_ = 2.74 ± 0.01 without the inducer and 2.76 ± 0.16 with inducer) was chosen for the further study here due to its shorter lag phase compared to the other engineered strains (Fig. 2).

All *pfl B*s tested showed improved maximum OD compared to *C. ljungdahlii*[empty], both in the presence and absence of Theophylline (Fig. 2). It seems that *pfl A* (Awo_c14940) can interact and function with all *pfl B*s from *A. woodii*. The two *pfl A* (Awo_c14910 and Awo_c14940) from *A. woodii* shares 47 % identities in protein sequences. Other *pfl B*s that might be able to interact better with *pfl A* (Awo_c14910) could not be cloned. Nonetheless, heterologous expression of *pfl B1* (Awo_c14920) showed to be the most effective with *pfl A* (Awo_c14940) (Fig. 2). Moon et al., showed that *pfl B1* (Awo_c14920) is the most abundant transcript and upregulated *pfl B* among *pfl B*s in *A. woodii* under formatotrophic condition, as well as *pfl A1* (Awo_c14910) (23). Another plausible explanation of *pfl B1* to be the most effective is that “leaky” expression of other *pfl B*s from the Theophylline-responsive riboswitch is also toxic to *C. ljungdahlii* making the lag phase longer.

The initial design of Theophylline riboswitch had a distinguishable background expression as usual natural bacterial promoters (34, 43). Previous studies using Theophylline-responsive riboswitches screened riboswitch sequences for showing low background expression compared to the parental design (29, 34). However, it seems that there is still a marginal background expression from the Theophylline-responsive riboswitch used, considering that PFL expressing *C. ljungdahlii* showed improvement in cell growth in the absence of Theophylline (Fig. 2). Nevertheless, in this study, *C. ljungdahlii* [B1A2] showed the same lag phase duration and similar maximum OD regardless of the presence or absence of the inducer, while cell growth was slightly lower in the presence (*p*-value = 0.16) than in the absence of the inducer (Fig. 2). Therefore, we decided not to add Theophylline to avoid the metabolic burden coming from the addition of the inducer for PFL expression.

### 3.4 The effect of different sodium formate concentrations on the engineered strain

When 80 mM of sodium formate was used for the cultivation of the engineered the strains, all the engineered strains showed 2 – 4 days lag phases. Different concentrations of sodium formate were tested to see if the amount of sodium formate has an impact on the lag phase.

When grown in 80 mM sodium formate, *C. ljungdahlii* [empty] showed around 2 days of lag phase with slower growth rate (*μ* = 0.058 ± 0.003 h^-1^) than in 0 mM (*μ* = 0.099 ± 0.007 h^-1^, *p*-value = 0.004) and in 40 mM (*μ* = 0.090 ± 0.003 h^-1^, *p*-value = 0.0004) of sodium formate (Fig. 3A). *C. ljungdahlii* [empty] in 0 mM and 40 mM of sodium formate did not show such long lag phase (Fig. 3A).

The maximum OD of *C. ljungdahlii* [empty] was slightly increased with sodium formate (OD_max, 40 mM_ =1.76 ± 0.08 with *p*-value = 0.013 and OD_max, 80 mM_ =1.80 ± 0.15 with *p*-value = 0.045) from without sodium formate (OD_max, 0 mM_ =1.47 ± 0.09) (Fig. 3A). Addition of sodium formate in the medium increased acetate production of *C. ljungdahlii* [empty] from 155 ± 6 mM to 178 ± 15 mM (with 40 mM sodium formate) and 206 ± 23 mM (with 80 mM sodium formate) (Fig. S2). On the other hand, when sodium formate was added to the growth medium with *C. ljungdahlii* [B1A2], the maximum OD was greatly improved from OD_max, 0 mM_ =1.86 ± 0.17 to OD_max, 40 mM_ =2.45 ± 0.21 and OD_max, 80 mM_ =3.07 ± 0.25 (Fig. 3B). However, 80 mM of sodium formate also decreased the growth rate of *C. ljungdahlii* [B1A2] to *μ* = 0.077 ± 0.004 h^-1^, compared to 0 mM (*μ* = 0.096 ± 0.001 h^-1^, *p*-value = 0.008) and 40 mM (*μ* = 0.088 ± 0.008 h^-1^, *p*-value = 0.151) of sodium formate (Fig. 3A and B). The amount of acetate or other metabolites produced remained the same whether or not *pfl* from *A. woodii* was expressed (Fig. S2 and 3).

Formate can be used in *C. ljungdahlii* as a substrate for formate dehydrogenase and as an intermediate chemical in the Wood-Ljungdahl pathway (2, 44). Considering that most formate was consumed in the early log phases, formate consumption was not related with cell growth of the control and engineered strain (Fig. S2 and S3). On the one hand, the maximum OD of *C. ljungdahlii* [empty] did not change with the concentration of sodium formate added. On the other hand, the maximum ODs and the amount of acetate produced by *C. ljungdahlii* [B1A2] increased with the amount of sodium formate added, as well as the amount of acetate produced from *C. ljungdahlii* [empty] using different formate concentrations. Moon et al., showed that formate consumption by *A. woodii* leads to pH increase (23). Therefore, it migh be that pH increase resulting from formate consumption contribute to increased acetate production. Also, the cell growth of *C. ljungdahlii* [B1A2] seemed to increase proportionally with acetate production, as the rise in medium pH due to formate consumption was counterbalanced by acidification from acetate production (Fig. S2). Nonetheless, the maximum OD of *C. ljungdahlii* [B1A2] was increased compared to the one of *C. ljungdahlii* [empty] even without supplementation of sodium formate (*p*-value = 0.04). Therefore, it can be concluded that heterologous expression of PFL from *A. woodii* has a positive impact on the cell growth of *C. ljungdahlii* under H_2_:CO_2_ condition.

**Figure 3.**
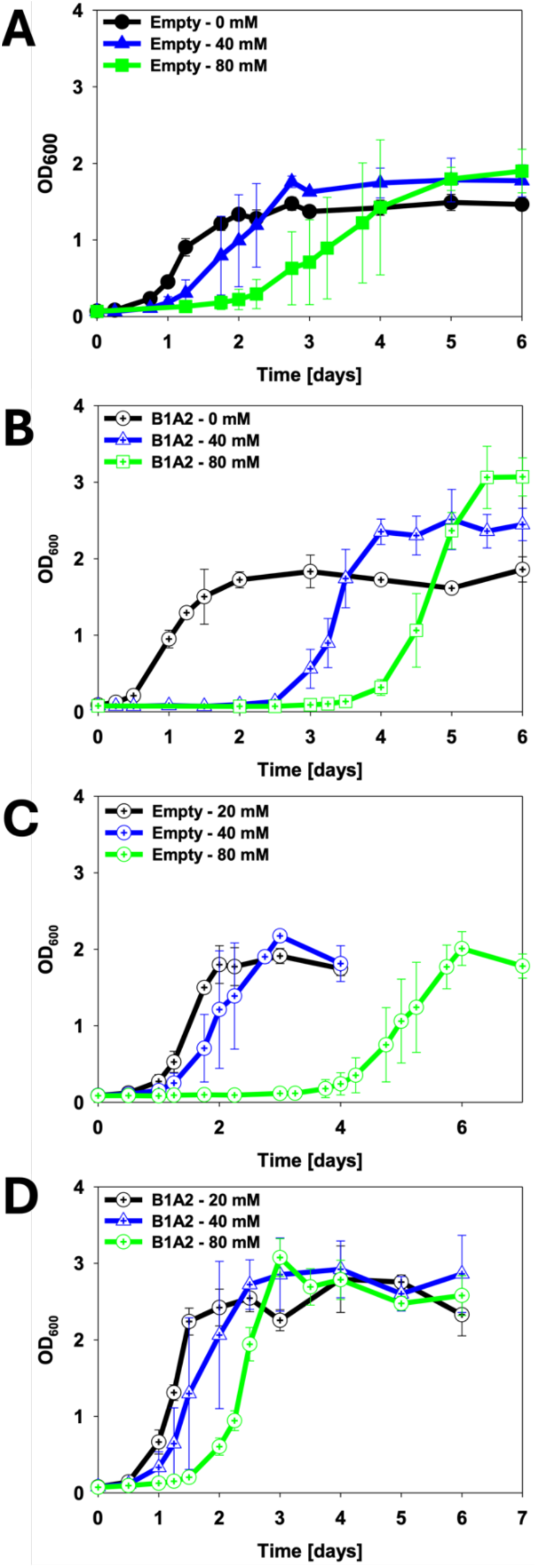
Effect of different sodium formate concentrations on growth of A) C. ljungdahlii [empty], B) C.ljungdahlii [B1A2], and C) 80 mM sodium formate-adapted C. ljungdahlii [empty] and D) 80 mM sodium formate-adapted C. ljungdahlii [B1A2] under H_2_:CO_2_ condition. Error bars indicate standard deviation (n=3).

### 3.5 The effect of adaptation on sodium formate

The long lag phase observed from the cultivation of the engineered strains in 40 mM and 80 mM sodium formate might be detrimental to *C. ljungdahlii* in a BES in which cell growth of *C. ljungdahlii* is inhibited (Fig. 3A and B) (3, 10). Therefore, the effect of adaptation of the engineered *C. ljungdahlii* strains on 80 mM sodium formate was investigated to see if the lag phase can be shortened. *C. ljungdahlii* [empty] and *C. ljungdahlii* [B1A2] cultivated under 80 mM formate condition were re-inoculated in different concentrations (20, 40, and 80 mM) of sodium formate.

Even after the adaptation of *C. ljungdahlii* [empty] on 80 mM sodium formate, neither the time for lag phase shortened nor the growth rates increased (*μ* = 0.096 ± 0.004 h^-1^, 0.092 ± 0.004 h^-1^, and 0.062 ± 0.009 h^-1^ at 0, 40, and 80 mM formate, respectively) (Fig. 3A and C). However, the maximum OD of *C. ljungdahlii* [empty] was increased to 1.91 ± 0.10 (20 mM), 2.17 ± 0.06 (40 mM), and 2.01 ± 0.22 (80 mM) after adaptation compared to 1.47 ± 0.09 (0 mM), 1.76 ± 0.08 (40 mM), and 1.80 ± 0.15 (80 mM) before adaptation. The amount of acetate produced was also increased compared to before adaptation (Fig. S3). Meanwhile, when *pfl* from *A. woodii* was heterologously expressed in *C. ljungdahlii*, not only was the 3 days lag phase shortened to 1 day in 80 mM sodium formate, but the growth rates in all sodium formate concentrations tested were also increased compared to before adaptation (*μ* = 0.124 ± 0.013 h^-1^, 0.113 ± 0.005 h^-1^, and 0.092 ± 0.002 h^-1^ at 20, 40 and 80 mM, respectively) (Fig. 3B and D).

The amount of acetate produced was also increased compared to before adaptation (Fig. S3 and 4). Compared to the control strain, *C. ljungdahlii* [B1A2] showed significant improvement in cell growth rate, maximum OD, and the time for the lag phase. It has been shown that *Escherichia coli* strains engineered for formate assimilation could be adaptively evolved for better utilization of formate (45, 46). Even though the amount of acetate produced and the maximum OD of *C. ljungdahlii* [empty] also increased with adaptation to sodium formate, *C. ljungdahlii* [B1A2] seemed to be more receptive for adaptation to efficient formate utilization for growth because of the heterologous PFL expression.

### 3.6 Bioelectrochemical reactor operation with the engineered strain

The engineered *C. ljungdahlii* strains showed significant improvement in cell growth under H_2_:CO_2_ condition with heterologous expression of *pfl* from *A. woodii*, supplementation of sodium formate, and adaptation on 80 mM sodium formate, compared to the control strain (Fig. 3). Therefore, the control and engineered strains were tested in a BES to see if the enhanced cell growth of *C. ljungdahlii* could be observed also in a BES. It was first tested with initial pH at pH 5.7. However, the pH increase in the cathode chamber hindered the cell growth, which masks the true effect of *pfl* from *A.woodii* expression in a BES (data not shown). Therefore, the experiment in the BES was conducted with the initial pH 5.0 for the engineered strains to perform its best as possible (3). Since *C. ljungdahlii* could grow only on 10 and 20 mM sodium formate with initial pH 5.0, previous flask experiments were not revisited for the BES reactor operation (Table 2). Furthermore, the amount of formate used for the BES experiment was 20 mM in the cathode chamber since 20 mM sodium formate was the highest formate concentration that *C. ljungdahlii* can tolerate at initial pH 5.0 (Table 2).

*C. ljungdahlii* [empty] showed very long lag phase in a BES for the first 5 days, and the OD only increased slightly to OD 0.10 ± 0.06 on day 12, after which the OD began to decline, indicating the onset of the death phase (Fig. 4A).

Acetate was produced to 10.2 ± 2.5 mM on day 13. All added formate was consumed by day 11 after which it was produced again, reaching 10.1 mM by day 20, likely through bioelectrochemical CO_2_ reduction following the cessation of acetogenesis and formate assimilation (Fig. 4B) (3, 7). *C. ljungdahlii* [empty] clearly struggled with long lag phase and poor growth in the BES (Fig. 4A). It seems that MES for CO_2_ reduction must be even harsher when harboring the foreign empty vector, compared to previous results with wild-type *C. ljungdahlii* showing OD 0.08 after 24 hours with no lag phase (3). It is speculated that the delayed cell growth might facilitate lactate production rather than cell growth (Fig. 4B).

**Figure 4.**
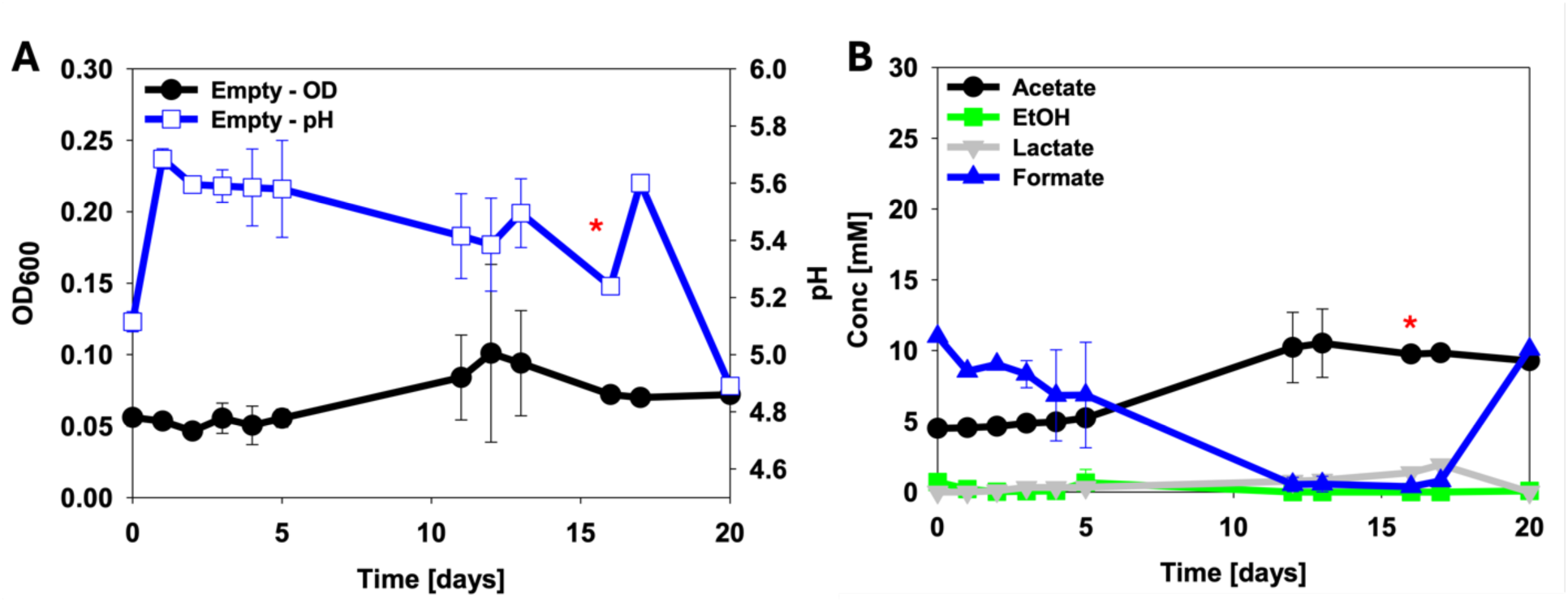
Time dependant A) OD, pH, and B) metabolites profile in a BES reactor using C. ljungdahlii [empty]. Metabolite concentrations are the mean of the cathode and anode chambers. Therefore, the initial formate concentration was aobut 10 mM because 20 mM sodium formate was added only to the cathode chamber. Error bars indicate standard deviation (n=2, the asterisk indicates n=1 after 15h).

**Figure 5.**
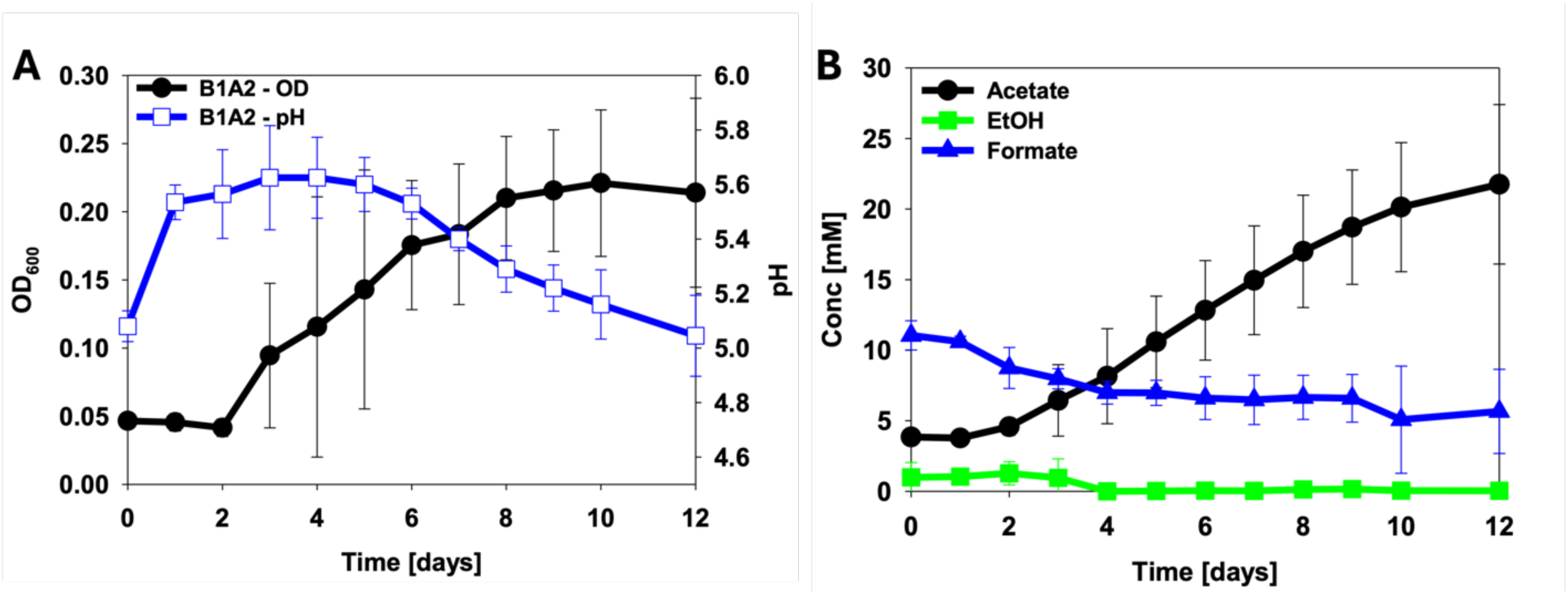
Time dependant A) OD, pH, and B) metabolites profiles in a BES reactor using C. ljungdahlii [B1A2]. Metabolite concentrations are the mean of the cathode and anode chambers. Therefore, the initial formate concentration was only about 10 mM because 20 mM sodium formate was added only to the cathode chamber. Error bars indicate standard deviation (n=2).

In contrast, *C. ljungdahlii* [B1A2] had two days of lag phase after which clear growth commenced, reaching the maximum OD 0.22 ± 0.05 on day 10 (Fig. 5A). Acetate was produced continuously to 21.8 ± 5.6 mM on day 12 (Fig. 5B). The formate concentration was maintained higher than 5 mM throughout the experiment (Fig. 5B). On the other hand, formate was fully consumed during the serum flask experiments whether or not PFL was heterologously expressed (Figs. S2-S5). Therefore, the sustained formate concentration may be due to bioelectrochemical CO_2_ reduction, with *C. ljungdahlii* [B1A2] steadily consuming formate, similar to *C. ljungdahlii* [empty] (3, 7). The extent of bioelectrochemical CO_2_ reduction to formate by *C. ljungdahlii* [B1A2] could be higher than for *C. ljungdahlii* [empty], as it produced more biomass than *C. ljungdahlii* [empty]. It could also be due to a more extensive CO_2_ fixation, stimulated by electrochemical H_2_ generation on the cathode and replacing part of the formate consumption.

High productivity is essential for MES to be commercially viable (47, 48). To be able to achieve high productivity, reaching higher current densities (100 mA cm^-2^ or higher) is often discussed by tuning operational parameters such as increasing salinity and CO_2_ solubility for faster reaction rate (48). However, cell growth of an acetogen in a BES, represented by the thickness of a biofilm on the cathode or by the OD, should be high enough to support such high current densities (48). Recently, Oriol et al., reported that up to – 28 ± 7 mA cm^-3^, which is comparable to rates observed in syngas fermentation, was achieved with a directed-flowthrough serpentine bioelectrochemical reactor using a mixed culture for a highly colonized biocathode (49). While utilizing mixed culture is essential for biofilm-driven MES (49), MES using a pure culture faces problems with low cell growth and thin biofilm formation (3, 10, 11). Therefore, a strategy for promoting cell growth in BES is necessary.

Cell growth of gas-fermenting *C. ljungdahlii*, as well as acetate production, was improved under H_2_:CO_2_ and MES conditions with PFL expression. This can be even more improved if PFL were integrated into the genome by genome editing techniques without the burden connected with the replication of foreign plasmids (29, 50). Also, heterologous co-expression of formyl-THF synthetase 2 (Awo_c08040) and formate transporter (Awo_c08050) from *A. woodii* might improve formate availability by *C. ljungdahlii* in a BES without leaving trace amount of residual formate (23).

*Clostridium* genus has been using strong constitutive promoters and has a limited number of inducible promoter systems. None of the inducible promoters has shown tight regulation without leaky expression (43). Therefore, more delicate expression systems are required to study this kind of toxic but beneficial genes. Even though there are several studies about promoter engineering and development of new promoter systems, it is challenging to develop a new strain without a tightly regulated promoter systems with various dynamic ranges (43, 51–54). In this study, we took advantage of “leaky” expression of the theophylline riboswitch to express PFL, that can be toxic when overexpressed. Also, *pfl A* was constitutively overexpressed from the plasmid in *C. ljungdahlii*. If other transcriptional inducible promoter systems were used in this situation, more balanced expression of *pfl B* and *pfl A* may be possible. This may potentially lead to further stimulation of growth and metabolic activity by elimination of detrimental effects of the heterologous gene expression (53, 54).

## 4. Conclusion

MES is an emerging way of renewable commodity chemical production. Utilizing an acetogen pure culture in a BES can offer immense opportunities to produce value-added chemicals by reassimilating geenhouse gas. Yet, the limited electron supply in a BES challenges the growth of acetogens. Carbon flux re-distribution to pyruvate synthesis, bypassing ferredoxin utilization, through PFL from *A. woodii* enabled significantly improved cell growth of *C. ljungdahlii* under H_2_:CO_2_ condition. Adaptive evolution on 80 mM sodium formate further improved the cell growth of the engineered *C. ljungdahlii* strain possessing PFL. The cell growth of the engineered *C. ljungdahlii* strain was also improved in a BES, resulting also in improved acetate production. These results will form a foundation for further strain development for MES as well as for gas fermentation technology.

## Acknowledgement

We thank Prof. Nigel Minton from the University of Nottingham, United Kingdom for kindly providing the pMTL80000 modular *Clostridium*-*E.coli* shuttle plasmids. This work was supported by the Swedish Energy Agency (grant number 46605-1) and the Area of Advance Energy, Chalmers University of Technology.

## References

1. Yishai O, Lindner SN, Gonzalez de la Cruz J, Tenenboim H, Bar-Even A. The formate bio-economy. Curr Opin Chem Biol. 2016;35:1–9.

2. Kopke M, Held C, Hujer S, Liesegang H, Wiezer A, Wollherr A, et al. Clostridium ljungdahlii represents a microbial production platform based on syngas. Proc Natl Acad Sci U S A. 2010;107(29):13087–92.

3. Im C, Valgepea K, Modin O, Nygård Y. Clostridium ljungdahlii as a biocatalyst in microbial electrosynthesis – Effect of culture conditions on product formation. Bioresour Technol Rep. 2022;19:101156.

4. Liu H, Song T, Fei K, Wang H, Xie J. Microbial electrosynthesis of organic chemicals from CO2 by Clostridium scatologenes ATCC 25775T. Bioresources and Bioprocessing. 2018;5(1).

5. Nevin KP, Hensley SA, Franks AE, Summers ZM, Ou J, Woodard TL, et al. Electrosynthesis of organic compounds from carbon dioxide is catalyzed by a diversity of acetogenic microorganisms. Appl Environ Microbiol. 2011;77(9):2882–6.

6. Nevin KP, Woodard TL, Franks AE, Summers ZM, Lovley DR. Microbial electrosynthesis: feeding microbes electricity to convert carbon dioxide and water to multicarbon extracellular organic compounds. mBio. 2010;1(2).

7. Im C, Kim M, Kim JR, Valgepea K, Modin O, Nygård Y, et al. Low electric current in a bioelectrochemical system facilitates ethanol production from CO using CO-enriched mixed culture. Frontiers in Microbiology. 2024;15.

8. Liew F, Martin ME, Tappel RC, Heijstra BD, Mihalcea C, Kopke M. Gas Fermentation-A Flexible Platform for Commercial Scale Production of Low-Carbon-Fuels and Chemicals from Waste and Renewable Feedstocks. Front Microbiol. 2016;7:694.

9. Marcellin E, Behrendorff JB, Nagaraju S, DeTissera S, Segovia S, Palfreyman RW, et al. Low carbon fuels and commodity chemicals from waste gases – systematic approach to understand energy metabolism in a model acetogen. Green Chem. 2016;18(10):3020–8.

10. Boto ST, Bardl B, Harnisch F, Rosenbaum MA. Microbial electrosynthesis with Clostridium ljungdahlii benefits from hydrogen electron mediation and permits a greater variety of products. Green Chem. 2023;25(11):4375–86.

11. Bajracharya S, ter Heijne A, Dominguez Benetton X, Vanbroekhoven K, Buisman CJ, Strik DP, et al. Carbon dioxide reduction by mixed and pure cultures in microbial electrosynthesis using an assembly of graphite felt and stainless steel as a cathode. Bioresour Technol. 2015;195:14–24.

12. Leang C, Ueki T, Nevin KP, Lovley DR. A genetic system for Clostridium ljungdahlii: a chassis for autotrophic production of biocommodities and a model homoacetogen. Appl Environ Microbiol. 2013;79(4):1102–9.

13. Han S, Gao X-y, Ying H-j, Zhou CC. NADH gene manipulation for advancing bioelectricity in Clostridium ljungdahlii microbial fuel cells. Green Chemistry. 2016;18(8):2473–8.

14. Klask CM, Kliem-Kuster N, Molitor B, Angenent LT. Nitrate Feed Improves Growth and Ethanol Production of *Clostridium ljungdahlii* With CO2 and H2, but Results in Stochastic Inhibition Events. Front Microbiol. 2020;11:724.

15. Zhu HF, Liu ZY, Zhou X, Yi JH, Lun ZM, Wang SN, et al. Energy Conservation and Carbon Flux Distribution During Fermentation of CO or H2/CO2 by *Clostridium ljungdahlii*. Front Microbiol. 2020;11:416.

16. Heffernan JK, Valgepea K, de Souza Pinto Lemgruber R, Casini I, Plan M, Tappel R, et al. Enhancing CO2-Valorization Using *Clostridium autoethanogenum* for Sustainable Fuel and Chemicals Production. Front Bioeng Biotechnol. 2020;8:204.

17. Tremblay PL, Zhang T, Dar SA, Leang C, Lovley DR. The Rnf complex of *Clostridium ljungdahlii* is a proton-translocating ferredoxin:NAD+ oxidoreductase essential for autotrophic growth. mBio. 2012;4(1):e00406–12.

18. Wu C, Lo J, Urban C, Gao X, Yang B, Humphreys J, et al. Acetyl-CoA synthesis through a bicyclic carbon-fixing pathway in gas-fermenting bacteria. Nature Synthesis. 2022;1(8):615–25.

19. Gupta R, Gupta N. Fundamentals of Bacterial Physiology and Metabolism 2021.

20. Aklujkar M, Leang C, Shrestha PM, Shrestha M, Lovley DR. Transcriptomic profiles of *Clostridium ljungdahlii* during lithotrophic growth with syngas or H2 and CO2 compared to organotrophic growth with fructose. Sci Rep. 2017;7(1):13135.

21. Claassens NJ. Reductive Glycine Pathway: A Versatile Route for One-Carbon Biotech. Trends Biotechnol. 2021;39(4):327–9.

22. Song Y, Lee JS, Shin J, Lee GM, Jin S, Kang S, et al. Functional cooperation of the glycine synthase-reductase and Wood-Ljungdahl pathways for autotrophic growth of *Clostridium drakei*. Proc Natl Acad Sci U S A. 2020;117(13):7516–23.

23. Moon J, Donig J, Kramer S, Poehlein A, Daniel R, Muller V. Formate metabolism in the acetogenic bacterium *Acetobacterium woodii*. Environ Microbiol. 2021;23(8):4214–27.

24. Zelcbuch L, Lindner SN, Zegman Y, Vainberg Slutskin I, Antonovsky N, Gleizer S, et al. Pyruvate Formate-Lyase Enables Efficient Growth of *Escherichia coli* on Acetate and Formate. Biochemistry. 2016;55(17):2423–6.

25. Bar-Even A. Formate Assimilation: The Metabolic Architecture of Natural and Synthetic Pathways. Biochemistry. 2016;55(28):3851–63.

26. Wood JC, Gonzalez-Garcia RA, Daygon D, Talbo G, Plan MR, Marcellin E, et al. Characterisation of acetogen formatotrophic potential using *Eubacterium limosum*. Appl Microbiol Biotechnol. 2023.

27. Neuendorf CS, Vignolle GA, Derntl C, Tomin T, Novak K, Mach RL, et al. A quantitative metabolic analysis reveals *Acetobacterium woodii* as a flexible and robust host for formate-based bioproduction. Metab Eng. 2021;68:68–85.

28. Heap JT, Pennington OJ, Cartman ST, Minton NP. A modular system for *Clostridium* shuttle plasmids. J Microbiol Methods. 2009;78(1):79–85.

29. Canadas IC, Groothuis D, Zygouropoulou M, Rodrigues R, Minton NP. RiboCas: A Universal CRISPR-Based Editing Tool for *Clostridium*. ACS Synth Biol. 2019;8(6):1379–90.

30. Woods C, Humphreys CM, Rodrigues RM, Ingle P, Rowe P, Henstra AM, et al. A novel conjugal donor strain for improved DNA transfer into *Clostridium* spp. Anaerobe. 2019;59:184–91.

31. Liew F, Henstra AM, Kpke M, Winzer K, Simpson SD, Minton NP. Metabolic engineering of *Clostridium autoethanogenum* for selective alcohol production. Metab Eng. 2017;40:104–14.

32. Kirkpatrick C, Maurer LM, Oyelakin NE, Yoncheva YN, Maurer R, Slonczewski JL. Acetate and formate stress: opposite responses in the proteome of *Escherichia coli*. J Bacteriol. 2001;183(21):6466–77.

33. Warnecke T, Gill RT. Organic acid toxicity, tolerance, and production in *Escherichia coli* biorefining applications. Microb Cell Fact. 2005;4:25.

34. Lynch SA, Desai SK, Sajja HK, Gallivan JP. A high-throughput screen for synthetic riboswitches reveals mechanistic insights into their function. Chem Biol. 2007;14(2):173–84.

35. Nagaraju S, Davies NK, Walker DJ, Kopke M, Simpson SD. Genome editing of *Clostridium autoethanogenum* using CRISPR/Cas9. Biotechnol Biofuels. 2016;9:219.

36. Cui W, Han L, Cheng J, Liu Z, Zhou L, Guo J, et al. Engineering an inducible gene expression system for *Bacillus subtilis* from a strong constitutive promoter and a theophylline-activated synthetic riboswitch. Microb Cell Fact. 2016;15(1):199.

37. Rudolph MM, Vockenhuber MP, Suess B. Conditional control of gene expression by synthetic riboswitches in *Streptomyces coelicolor*. Methods Enzymol. 2015;550:283–99.

38. Nakahira Y, Ogawa A, Asano H, Oyama T, Tozawa Y. Theophylline-dependent riboswitch as a novel genetic tool for strict regulation of protein expression in Cyanobacterium *Synechococcus elongatus* PCC 7942. Plant Cell Physiol. 2013;54(10):1724–35.

39. Dwidar M, Yokobayashi Y. Controlling Bdellovibrio bacteriovorus Gene Expression and Predation Using Synthetic Riboswitches. ACS Synth Biol. 2017;6(11):2035–41.

40. Ohbayashi R, Akai H, Yoshikawa H, Hess WR, Watanabe S. A tightly inducible riboswitch system in *Synechocystis* sp. PCC 6803. J Gen Appl Microbiol. 2016;62(3):154-9.

41. Topp S, Reynoso CM, Seeliger JC, Goldlust IS, Desai SK, Murat D, et al. Synthetic riboswitches that induce gene expression in diverse bacterial species. Appl Environ Microbiol. 2010;76(23):7881–4.

42. Crain AV, Broderick JB. Pyruvate formate-lyase and its activation by pyruvate formate-lyase activating enzyme. J Biol Chem. 2014;289(9):5723–9.

43. Joseph RC, Kim NM, Sandoval NR. Recent Developments of the Synthetic Biology Toolkit for *Clostridium*. Front Microbiol. 2018;9:154.

44. Moon M, Park GW, Lee J-P, Lee J-S, Min K. Recombinant expression and characterization of formate dehydrogenase from *Clostridium ljungdahlii* (ClFDH) as CO2 reductase for converting CO2 to formate. Journal of CO2 Utilization. 2022;57.

45. Kim SJ, Yoon J, Im DK, Kim YH, Oh MK. Adaptively evolved *Escherichia coli* for improved ability of formate utilization as a carbon source in sugar-free conditions. Biotechnol Biofuels. 2019;12:207.

46. Delmas VA, Perchat N, Monet O, Foure M, Darii E, Roche D, et al. Genetic and biocatalytic basis of formate dependent growth of *Escherichia coli* strains evolved in continuous culture. Metab Eng. 2022;72:200–14.

47. Claassens NJ, Cotton CAR, Kopljar D, Bar-Even A. Making quantitative sense of electromicrobial production. Nat Catal. 2019;2(5):437–47.

48. Jourdin L, Burdyny T. Microbial Electrosynthesis: Where Do We Go from Here? Trends Biotechnol. 2021;39(4):359–69.

49. Cabau-Peinado O, Winkelhorst M, Stroek R, de Kat Angelino R, Straathof AJJ, Masania K, et al. Microbial electrosynthesis from CO2 reaches productivity of syngas and chain elongation fermentations. Trends Biotechnol. 2024.

50. Philipps G, de Vries S, Jennewein S. Development of a metabolic pathway transfer and genomic integration system for the syngas-fermenting bacterium *Clostridium ljungdahlii*. Biotechnol Biofuels. 2019;12:112.

51. Mordaka PM, Heap JT. Stringency of Synthetic Promoter Sequences in *Clostridium* Revealed and Circumvented by Tuning Promoter Library Mutation Rates. ACS Synth Biol. 2018;7(2):672–81.

52. Yang G, Jia D, Jin L, Jiang Y, Wang Y, Jiang W, et al. Rapid Generation of Universal Synthetic Promoters for Controlled Gene Expression in Both Gas-Fermenting and Saccharolytic *Clostridium* Species. ACS Synth Biol. 2017;6(9):1672–8.

53. Banerjee A, Leang C, Ueki T, Nevin KP, Lovley DR. Lactose-inducible system for metabolic engineering of *Clostridium ljungdahlii*. Appl Environ Microbiol. 2014;80(8):2410–6.

54. Hartman AH, Liu H, Melville SB. Construction and characterization of a lactose-inducible promoter system for controlled gene expression in *Clostridium perfringens*. Appl Environ Microbiol. 2011;77(2):471–8.

