## Supplementary information for "Heterologous expression of pyruvate formate lyase enhances cell growth of *Clostridium ljungdahlii* during microbial electrosynthesis"

### Supplymentary information

**Figure S1** Time-dependent indiviual OD profiles of different *pfl B* and *pfl A* combinations A)without and B) with addition of 1 mM of theophiline supplementation as the translational inducer for the riboswitch. Filled symbols indicates without addition of the inducer, and non-filled symboles indicates with addition of the inducer.

**Figure S2** The effect of different sodium formate concentrations on A) growth of *C. ljungdahlii* [empty]. Metabolites profile of *C. ljungdahlii* [empty] when B) 20 mM, C) 40 mM, and D) 80 mM sodium formate were added

**Figure S3** The effect of different sodium formate concentrations on A) growth of *C. ljungdahlii* [B1A2]. Metabolites profile of *C. ljungdahlii* [B1A2] when B) 20 mM, C) 40 mM, and D) 80 mM sodium formate were added

**Figure S4** The effect of different sodium formate concentrations on A) growth of *C. ljungdahlii* [empty] after adaptation on 80 mM sodium formate. Metabolites profile of 80 mM sodium formate adapted *C. ljungdahlii* [empty] when B) 20 mM, C) 40 mM, and D) 80 mM sodium formate were added

**Figure S5** The effect of different sodium formate concentrations on A) growth of *C. ljungdahlii* [B1A2] after adaptation on 80 mM sodium formate. Metabolites profile of 80 mM sodium formate adapted *C. ljungdahlii* [B1A2] when B) 20 mM, C) 40 mM, and D) 80 mM sodium formate were added

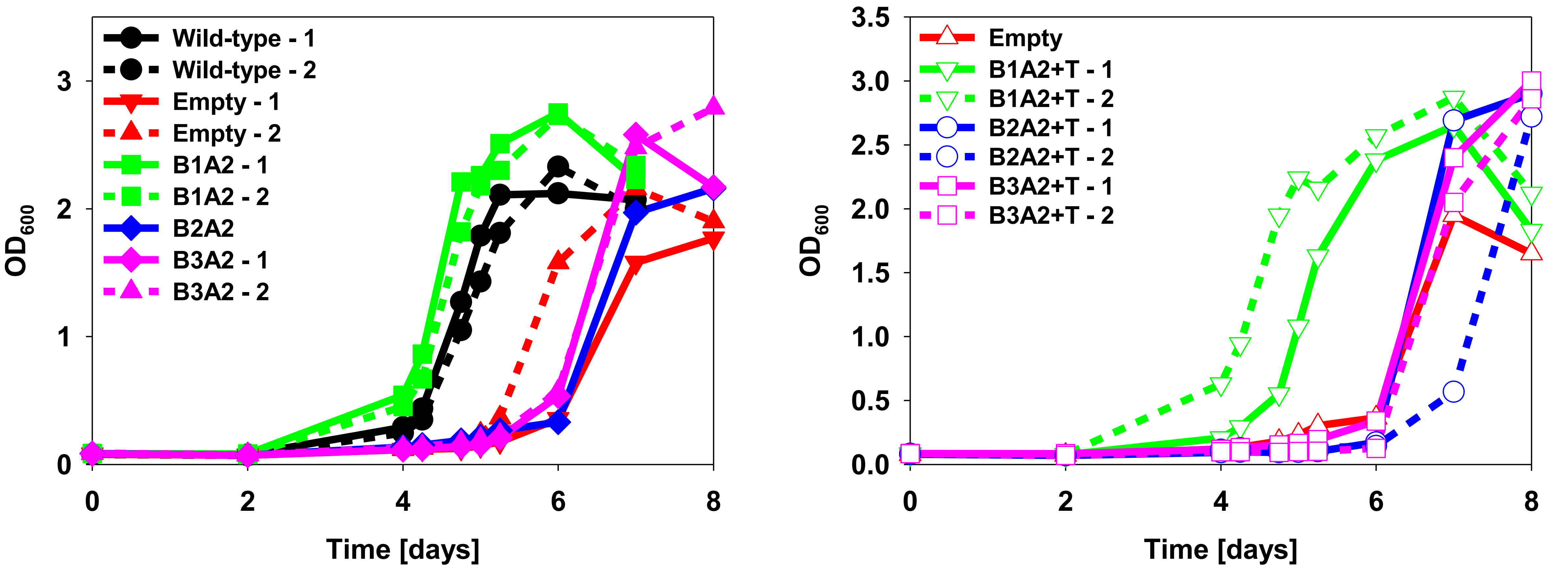


**Figure S1** Time-dependent indiviual OD profiles of different *pfl B* and *pfl A* combinations A)without and B) with addition of 1 mM of theophiline supplementation as the translational inducer for the riboswitch. Filled symbols indicates without addition of the inducer, and non-filled symboles indicates with addition of the inducer.

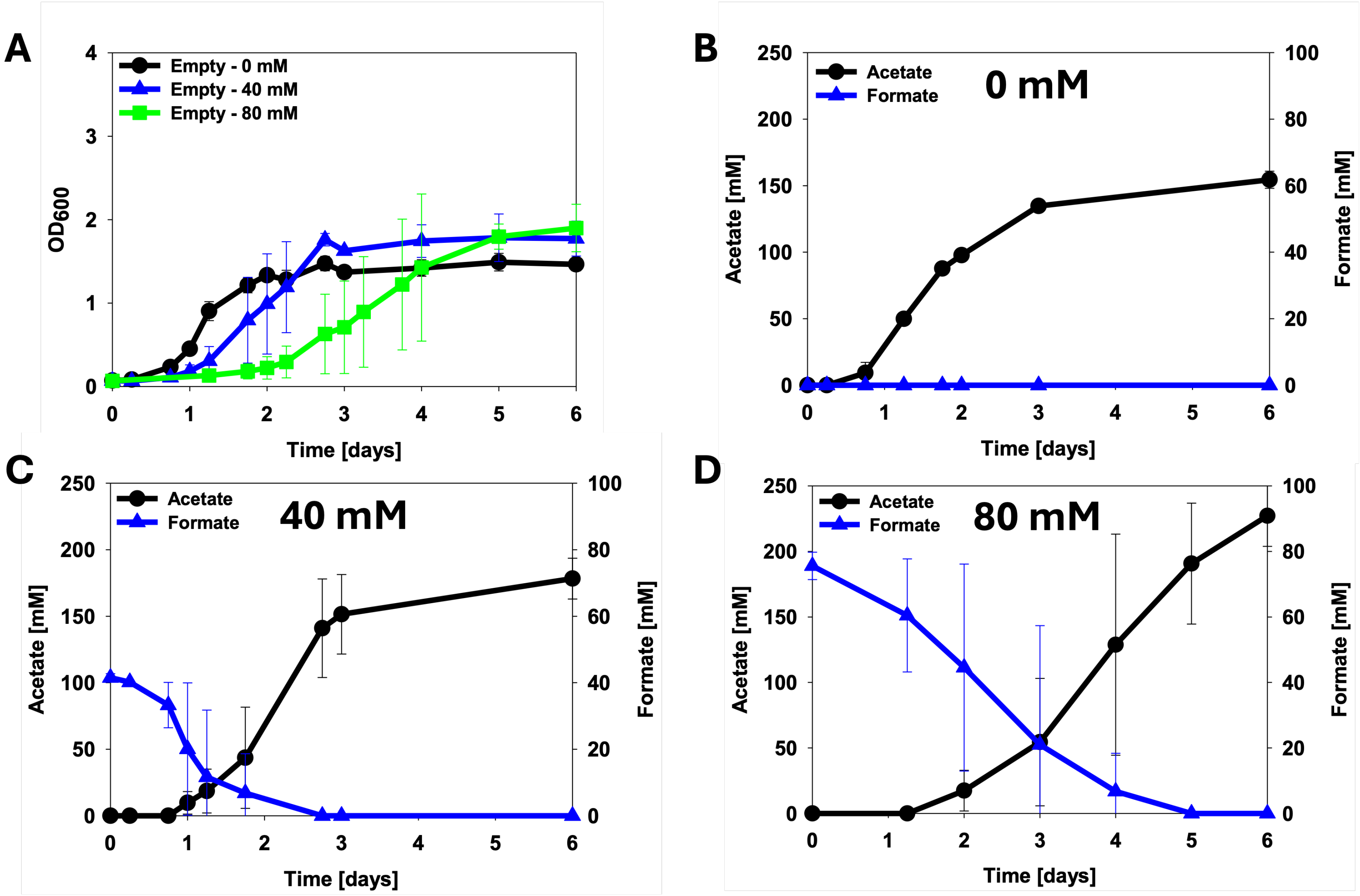


**Figure S2** The effect of different sodium formate concentrations on A) growth of *C. ljungdahlii* [empty]. Metabolites profile of *C. ljungdahlii* [empty] when B) 20 mM, C) 40 mM, and D) 80 mM sodium formate were added. Error bars indicate standard deviation (n=3).

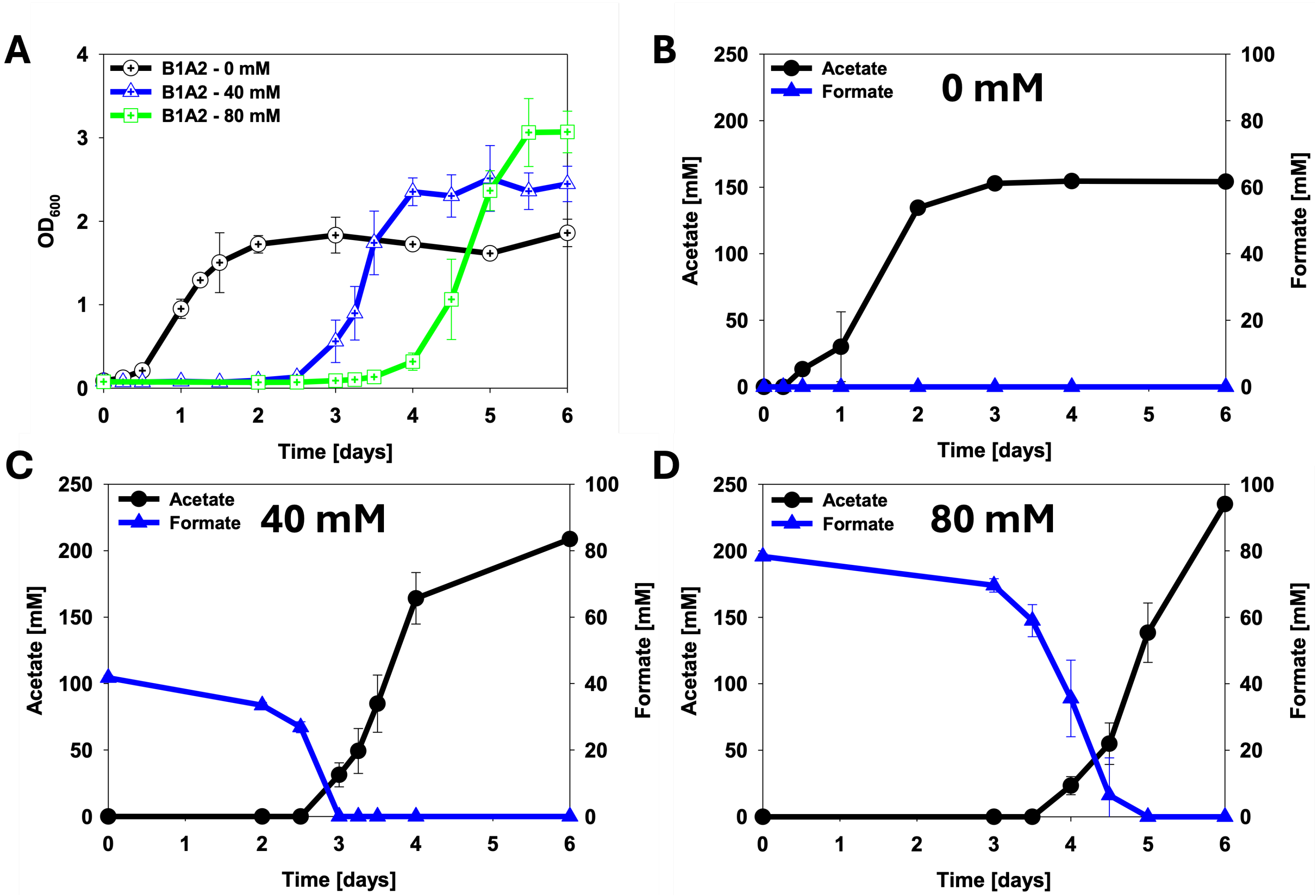


**Figure S3** The effect of different sodium formate concentrations on A) growth of *C. ljungdahlii* [B1A2]. Metabolites profile of *C. ljungdahlii* [B1A2] when B) 20 mM, C) 40 mM, and D) 80 mM sodium formate were added. Error bars indicate standard deviation (n=3).

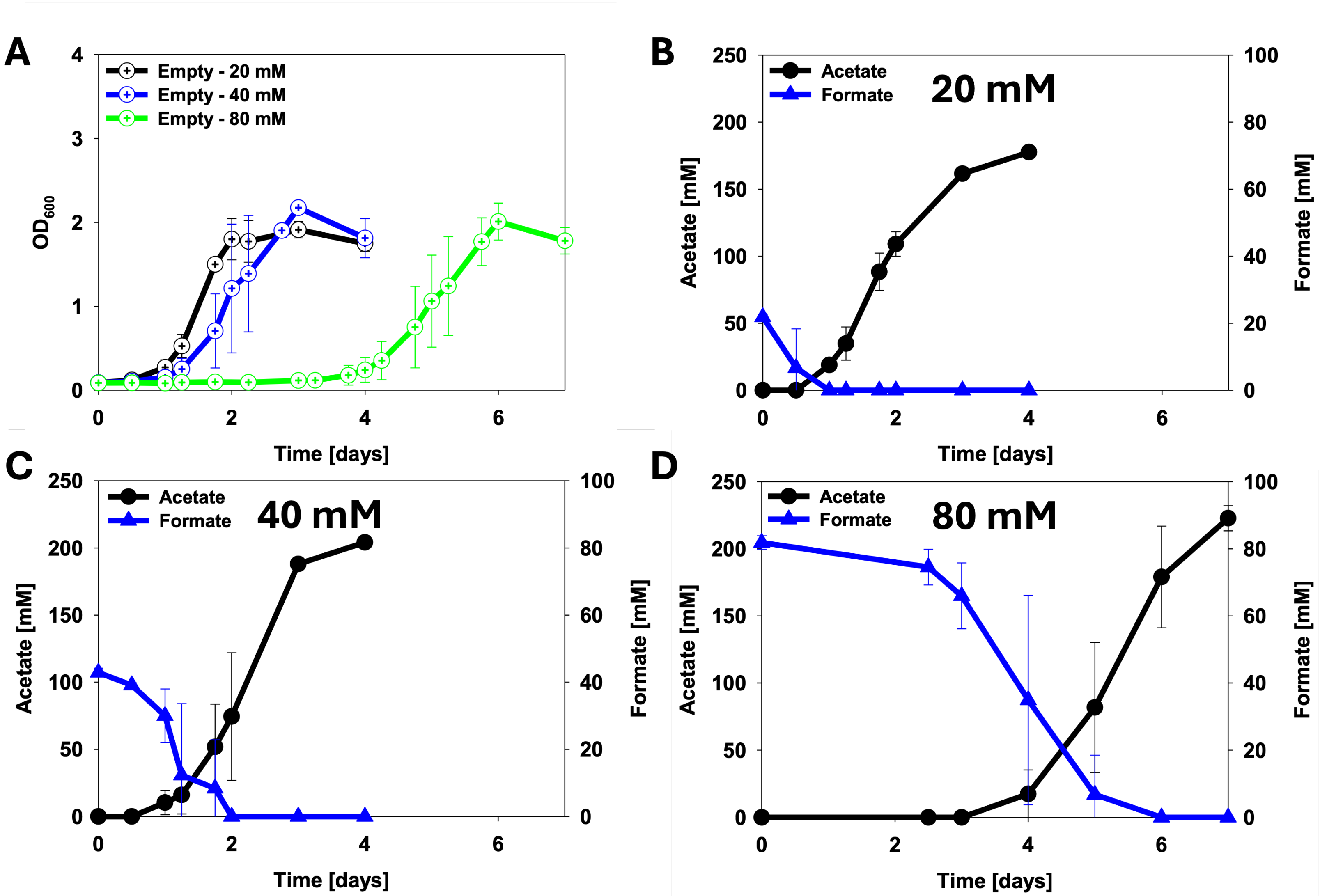


**Figure S4** The effect of different sodium formate concentrations on A) growth of *C. ljungdahlii* [empty] after adaptation on 80 mM sodium formate. Metabolites profile of 80 mM sodium formate adapted *C. ljungdahlii* [empty] when B) 20 mM, C) 40 mM, and D) 80 mM sodium formate were added. Error bars indicate standard deviation (n=3).

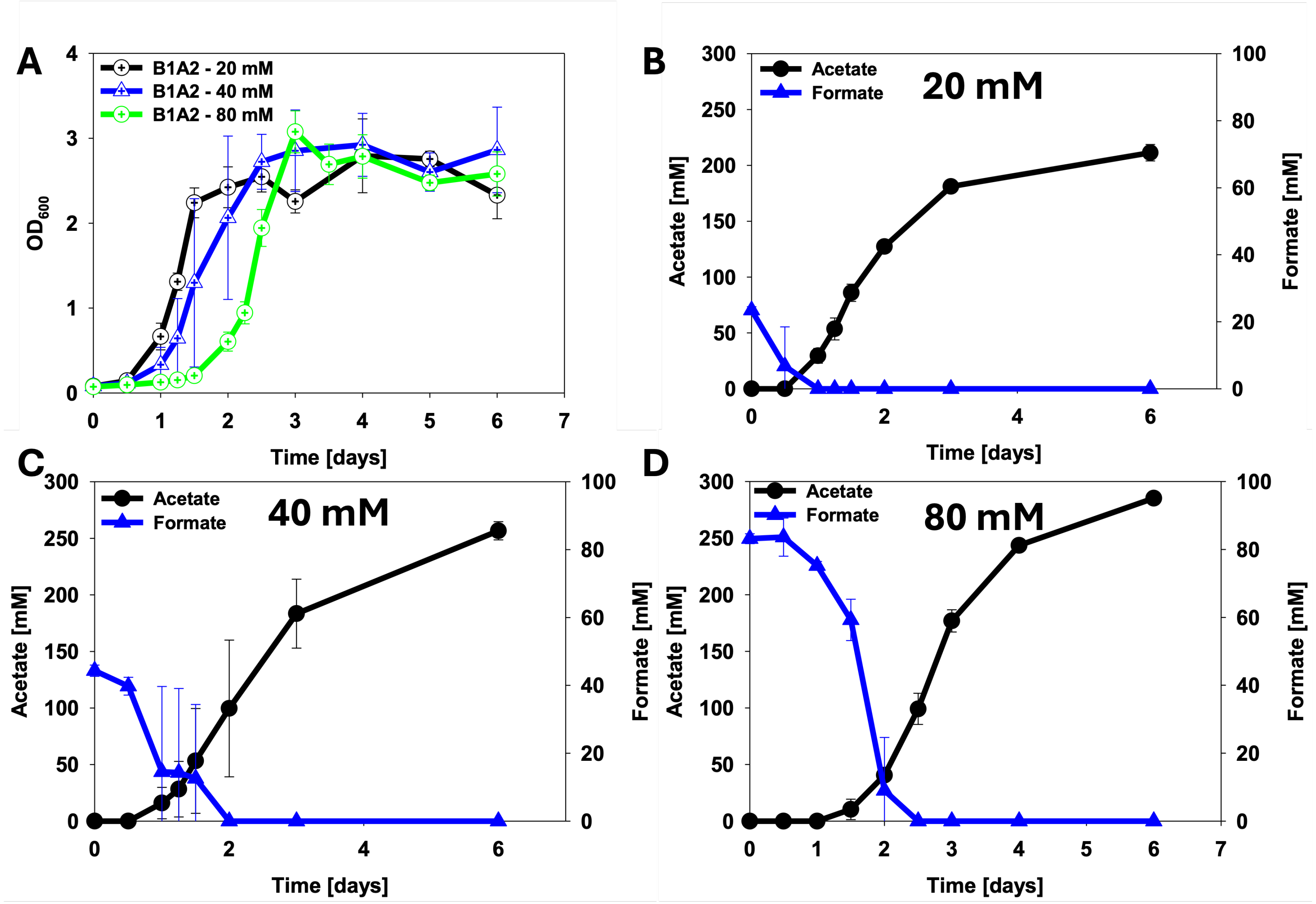


**Figure S5** The effect of different sodium formate concentrations on A) growth of *C. ljungdahlii* [B1A2] after adaptation on 80 mM sodium formate. Metabolites profile of 80 mM sodium formate adapted *C. ljungdahlii* [B1A2] when B) 20 mM, C) 40 mM, and D) 80 mM sodium formate were added. Error bars indicate standard deviation (n=3).
